# Importin-9 regulates chromosome segregation and packaging in *Drosophila* germ cells

**DOI:** 10.1101/2020.02.10.942326

**Authors:** Victor Palacios, Garrett C Kimble, Tina L. Tootle, Michael Buszczak

## Abstract

Germ cells undergo distinct nuclear processes as they differentiate into gametes. While these events must be coordinated to ensure proper maturation, the stage-specific transport of proteins in and out of germ cell nuclei remains incompletely understood.

Our efforts to genetically characterize *Drosophila* genes that exhibit enriched expression in germ cells led to the finding that loss of the highly-conserved Importin β/karyopherin family member Importin-9 (Ipo9) results in female and male sterility.

Immunofluorescence and fluorescent in situ hybridization (FISH) revealed that *Ipo9^KO^* mutants display chromosome condensation and segregation defects during meiosis. In addition, *Ipo9^KO^* mutant males form abnormally structured sperm and fail to properly exchange histones for protamines. Ipo9 physically interacts with proteasome proteins and *Ipo9* mutant males exhibit loss of nuclear ubiquitination and disruption of the nuclear localization of several proteasome components. Thus, Ipo9 coordinates the nuclear import of functionally related factors necessary for the completion of gametogenesis.

**Highlights:** *Drosophila* Importin-9 functions in female and male meiosis

Loss of Importin-9 disrupts the histone to protamine transition during spermiogenesis Importin-9 physically interacts with components of the proteasome

Importin-9 is required for the efficient nuclear transport of proteasome proteins during spermiogenesis

## Introduction

Subcellular compartmentalization allows for complex modes of gene regulation in eukaryotic cells. The regulated and active transport of macromolecules between different compartments promotes cellular homeostasis and often drives differentiation. Transport of molecules from the cytoplasm to the nucleus depends on a family of proteins called karyopherins, as known as importins (Cagatay and Chook, 2018; Chook and Blobel, 2001). The karyopherin superfamily of transporters consists of importin α and importin β sub-groups. All the proteins within this karyopherin superfamily share tandem huntingtin, elongation factor 3, protein phosphatase 2A and mechanistic target of rapamycin (HEAT) repeats. These repeats allow these proteins to bind to various cargo proteins, which often, but not always, contain a nuclear localization signal within their peptide sequence. Karyopherins then transport these cargoes into the nucleus through nuclear pores.

Another key component of the transport machinery is the small GTPase Ran (Cautain et al., 2015). Cytoplasmic Ran is typically maintained in a GDP-bound state, while nuclear Ran binds GTP. This concentration gradient of GDP and GTP bound Ran provides a directional cue for transport of proteins between the cytoplasm and nucleus. Once importins enter the nucleus, high affinity interactions with RanGTP cause karyopherins to release their cargoes and recycle back to the cytoplasm.

Accumulating evidence suggests that β-karyopherins do not simply function as constitutive and redundant housekeeping proteins. Interactions between different β-karyopherins with specific cargoes depends not only on their overlapping expression patterns in time and space, but also on clear differences in the affinities of the physical interactions (Gontan et al., 2009; Kimura and Imamoto, 2014; Major et al., 2011; Plafker and Macara, 2002; Quan et al., 2008). For example, histones can bind to multiple β-karyopherins, but their affinities vary. For example, Kapβ2 and Imp5 exhibit very strong affinity for Histone H3, while Impβ, Imp4, Imp7, Imp9 and Imp*α* display weaker interactions (Soniat et al., 2016). Additionally, a previous study identified a group 468 cargoes for 12 β-karyopherins (Kimura et al., 2017). Three hundred and thirty two of these cargoes were unique to one β-karyopherin family member, suggesting a division of function amongst these transporters. Several β-karyopherin family members have been associated with specific diseases. Accumulating evidence shows that β-karyopherins are overexpressed in multiple tumors including melanoma, pancreatic, breast, colon, gastric, prostate, esophageal, lung cancer, and lymphomas (Fujii et al., 2018; Turner et al., 2012). Additionally, specific karyopherin-β proteins such as exportin-1 have been implicated in drug resistance in cancer (Mahipal and Malafa, 2016; Turner et al., 2014; Turner et al., 2012).

Many importins exhibit enriched expression in gonads and are functionally required during different stages of spermatogenesis and oogenesis across many species, including *Drosophila*. *Drosophila* ovaries are organized into discrete units called ovarioles, which contain a series of sequentially developing egg chambers. Each egg chamber is comprised of 16 germ cells, 15 nurse cells and one oocyte, surrounded by a layer of somatic follicle cells. The initiation of meiosis occurs early in oogenesis, marked by the formation of the synaptonemal complex (SC) and the generation of the programmed double strand breaks. After these first events, oocytes remain arrested in prophase 1 of meiosis until Stage 12, followed by prometaphase 1 at Stage 13 and metaphase 1 at Stage 14 (Hughes et al., 2018).

The *Drosophila* testis is structured as a closed-end coiled tube. At the tip of the testis, 10-14 germline stem cells (GSCs) surround a small cluster of somatic cells called the hub. GSCs typically divide asymmetrically to produce another GSC and a gonialblast. Gonialblasts become enveloped by two somatic cyst cells, which function in an analogous manner to the Sertoli cells of the mammalian testis (White-Cooper, 2010). The *Drosophila* gonialblast goes through four incomplete mitotic divisions to form an interconnected 16-cell spermatogonial cell cyst. Each spermatocyte within the cyst undergoes meiosis, resulting in the formation of cysts that contain 64 interconnected haploid cells. Immediately after the completion of meiosis, these cells enter the “onion stage”, which is marked by the appearance of a single hyperfused mitochondria, called the nebenkern, which appears layered in electron micrographs. Defects in meiosis can result in the appearance of fragmented nebenkern and alternations in the normal 1:1 ratio of nuclei and nebenkern.

Spermiogenesis is marked by nuclear elongation and chromatin reorganization. Nuclear elongation is dependent on microtubules from the basal body that associate with the nucleus (Fabian and Brill, 2012). Chromatin organization switches from a histone-based to protamine-based packaging in the late elongation stage (Rathke et al., 2007). During elongation, the nuclear envelope that is in contact with the basal body forms a cavity that fills with microtubules while the nucleus takes a “canoe” shape. During chromatin reorganization, histones are ubiquitinated by an unknown ubiquitin ligase and subsequently degraded by the proteasome at the later canoe stage, immediately before protamines are incorporated into the chromatin (Awe and Renkawitz-Pohl, 2010; Zhong and Belote, 2007). After histone removal, the transition like-protein (Tpl) is incorporated, which facilitates protamine incorporation (Rathke et al., 2007). In *Drosophila*, mature sperm contain Mst35Ba (protamine A), Mst35Bb (protamine B) and Mst77F (Rathke et al., 2010). Towards the end of spermiogenesis, sperm form their own membranes in a process called individualization (Fabian and Brill, 2012).

In *Drosophila*, mutants in several importins develop normally into adults, but exhibit various defects in fertility. Importin α2 mutant males exhibit a dramatic decrease in the formation of individualized and motile sperm, while mutant females produce small and deflated eggs with missing or fused dorsal appendages (Giarre et al., 2002; Mason et al., 2002). Similarly, mutations in Importin α1 also cause male and female sterility, marked by egg-laying defects in females and the formation of spermatocytes with abnormally large round nuclei in males and loss of Importin α3 leads to arrest of oogenesis (Mathe et al., 2000). The specific cargoes responsible for these phenotypes remain unknown.

Here, we report that null mutations in *Ipo9* cause disruption of chromosome segregation and condensation during meiosis in both female and male *Drosophila*. Previous results have shown that Ipo9 helps to traffic Actin, Histone H2A-H2B dimers and a variety of other factors into nuclei (Dopie et al., 2012; Kortvely et al., 2005; Matsumiya et al., 2013; Padavannil et al., 2019; Sokolova et al., 2018). We confirm that loss of *Drosophila* Ipo9 disrupts the accumulation of nuclear actin during oogenesis. In addition, we find Ipo9 promotes chromosome segregation during meiosis, and the exchange of histones for protamines during spermiogenesis. Biochemical experiments suggest that Ipo9 physically associates with proteasome components, and immunofluorescent studies show that loss of Ipo9 disrupts the normal trafficking of the proteasome into germ cell nuclei during spermiogenesis. Together, these data reveal new processes directly regulated by a specific nuclear transport factor during gametogenesis.

## Results

### Loss of Importin-9 results in sterility

We sought to genetically characterize genes that display enriched transcription within gonads based on publicly available modEncode RNA-seq data. According to these datasets, the Importin β/karyopherin family member Ranbp9 (CG5252) exhibits high levels of expression in both ovaries and testes relative to other tissues (http://flybase.org/reports/FBgn0037894). The name Ranbp9 has previously been used for genes that do not share extensive homology with one another across species. For example, the mammalian Ranbp9 gene shares closest homology to the *Drosophila* RanBPM gene, while mammalian Importin-9 (Ipo9) represents the closest homolog of *Drosophila* CG5252. Given these discrepancies, we have elected to call CG5252 Importin-9 (Ipo9) hereafter.

To determine whether Ipo9 functions during germ cell development in both females and males, we generated a molecular null mutation by replacing most of the Ipo9 coding sequence with a 3XP3-DsRed cassette using CRISPR/Cas9-mediated genomic engineering (Figure S1A). Independent isolates of this *Ipo9^KO^* mutation were homozygous viable, but exhibited female and male sterility. *Ipo9^KO^* homozygous females laid a comparable number eggs to *w^1118^* controls and their ovaries appeared grossly normal (Figure 1A-C). However, none of eggs from the *Ipo9* mutants hatched. Staining for α-Tubulin and DNA revealed loss of maternal *Ipo9* results in widespread mitotic catastrophes during the earliest embryonic divisions, marked by chromosome bridges, chromosome fragmentation, lack of chromosome condensation and an array of spindle defects (Figure 1D-E’).

**Fig 1.**
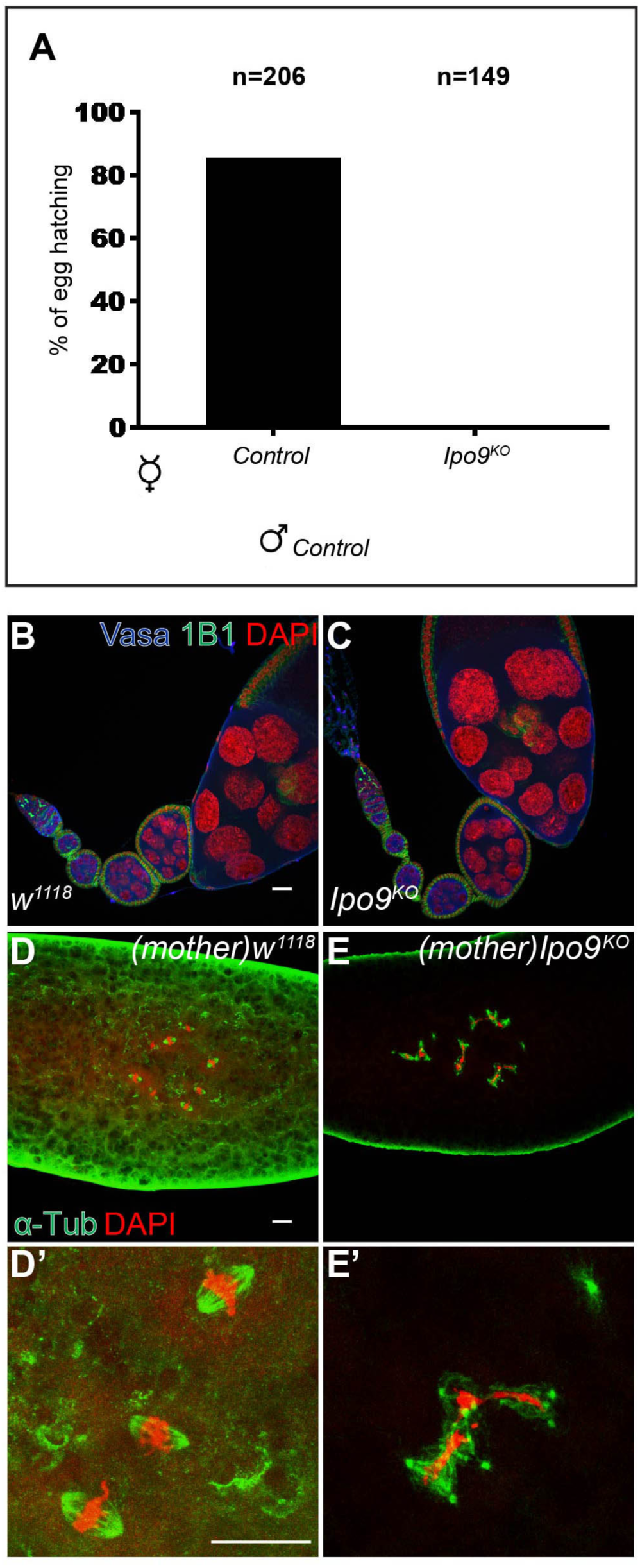
Embryos from *Ipo9^KO^* females show mitotic defects. (A) Percentage of eggs that hatch after 5 days of being laid by *w^1118^* or *Ipo9^KO^* females crosses with *w^1118^* males. (B-C) *Drosophila* ovarioles stained for VASA (blue), 1B1 (green) and DAPI (red). (B) *w^1118^* control and (C) *Ipo9^KO^* ovarioles. (D-D’) Embryos from *w^1118^* (control) and (E-E’) *Ipo9^KO^* females stained for α-Tub (green) and DAPI (red). Scale bars 20μm.

### The N-terminal beta-karyopherin domain is necessary for Importin-9 function

To verify that the female sterility of *Ipo9^KO^* homozygotes was caused by loss of *Ipo9*, and not disruption of another nearby gene, we used two methods: RNAi knockdown and cDNA rescue. Driving *Ipo9* specific RNAi using germ cell specific drivers resulted in the same female sterile phenotypes as the *Ipo9^KO^* mutant (Figure S1C-E). This result supports the idea that Ipo9 functions during gametogenesis. Moreover, these data indicate that *Ipo9* acts in a cell autonomous manner within germ cells to promote fertility.

To complement the RNAi knockdown experiments, we also attempted to rescue the *Ipo9^KO^* mutant with a full-length wild-type cDNA transgene (*UASp-Ipo9^FL^*). We made a second transgene (*UASp-Ipo9^ΔN^*), in which the N-terminal β-karyopherin domain was deleted (Figure 2A). This construct allowed us to test whether the *Ipo9* mutant phenotypes were caused by disruption of nuclear import of specific cargoes, as opposed potential transport independent functions. Both transgenes were expressed at similar levels but exhibited different rescuing activity and localization (Figure 2B-E’). While the Ipo9^FL^ HA-tagged transgenic protein was enriched around the nuclear envelop of nurse cells and appeared to enter the germinal vesicle within the oocyte, as expected, the Ipo9^ΔN^ protein did not, indicating that removal of this domain disrupted the ability of this protein to act as a nuclear importer (Figure 2D-E’). Driving the expression of the full-length transgene using *vasa-gal4* rescued the female sterile phenotypes, whereas the *Ipo9^ΔN^* construct did not (Figure 2C). Together, these results indicate that Ipo9-mediated nuclear trafficking is essential for normal gametogenesis in *Drosophila*.

**Fig 2.**
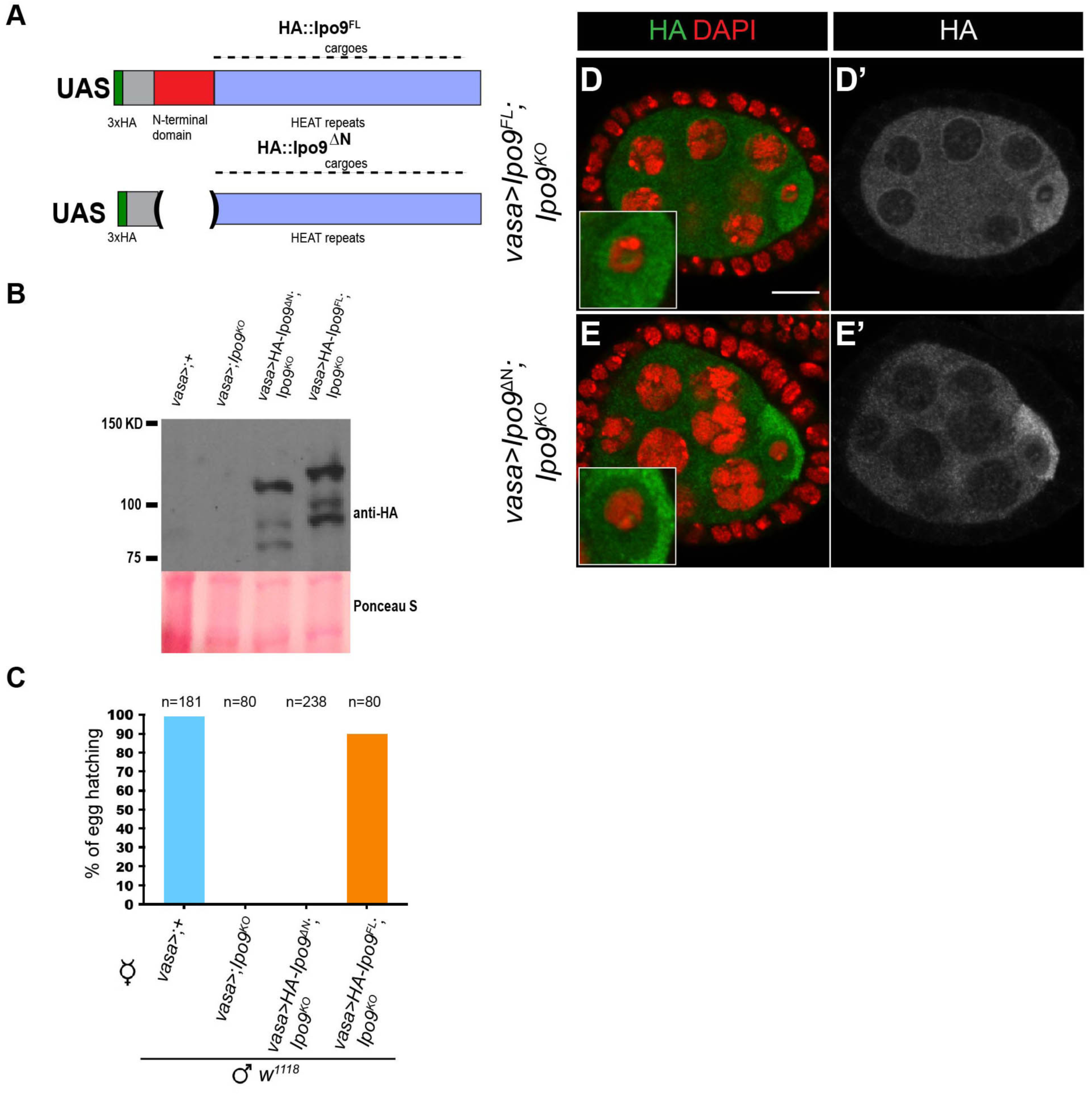
The N-terminal domain of Ipo9 is required for its function during gametogenesis. (A) Schematic of the 3XHA full length Ipo9 (Ipo9^FL^) and 3xHA DeltaN-Ipo9 (Ipo9^ΔN^) proteins. (B) Western blot from ovaries showing HA::Ipo9^ΔN^ and HA::Ipo9^FL^ expression. (C) Percentage of eggs that hatch after 5 days of being laid by *vasa-gal4>;+*, *vasa-gal4>;Ipo9^KO^*, *vasa-gal4>Ipo9^FL^;Ipo9^KO^* and *vasa-gal4>Ipo9*^ΔN^*;Ipo9^KO^* females crosses with *w^1118^* males. (D-E’) Stage 4-5 egg chambers stained for HA (green; grayscale) and DAPI (red) for these genotypes (D,D’) *vasa-gal4>Ipo9* ^FL^*;Ipo9^KO^* and (E, E’) *vasa-gal4>Ipo9*^ΔN^*;Ipo9^KO^* females. Scale bars 10μm.

### Loss of Ipo9 disrupts meiosis in females

Given that disruption of *Ipo9* leads to sterility in both females and males, we suspected that Ipo9 may play a role in meiosis. To characterize potential meiotic defects in *Ipo9* mutant females, we employed fluorescent in situ hybridization (FISH) using probes for the 359-bp repeat sequences near the X chromosome centromere and the AACAC(n) microsatellite repeats on the 2nd chromosome. In wild-type females, Stage 14 oocytes are arrested in metaphase phase I until ovulation and the chromatin of these oocytes appears as a single mass. FISH revealed that X-chromosome and 2nd chromosome pairs normally orient towards opposite poles (Figure 3A). However, *Ipo9* mutant oocytes tended to display mis-orientation of these chromosomes (Figure 3B-D). This phenotype was marked the appearance of individual X-chromosome and 2^nd^-chromosome spots in the middle of the nucleus or misorientation of all the chromosomes to one side of the nucleus, indicating that loss of *Ipo9* disrupts normal chromosome segregation patterns during meiosis. Examining Centrosome Identifier (CID), a centromere-specific histone H3 variant, in control and mutant meiotic nuclei provided additional evidence that loss of *Ipo9* results in chromosome mis-orientation during meiosis (Figure 3E-H). These defects were not correlated with disruption of the meiotic spindle, which appears largely normal in *Ipo9* mutant cells (Figure 3I-J’).

**Fig 3.**
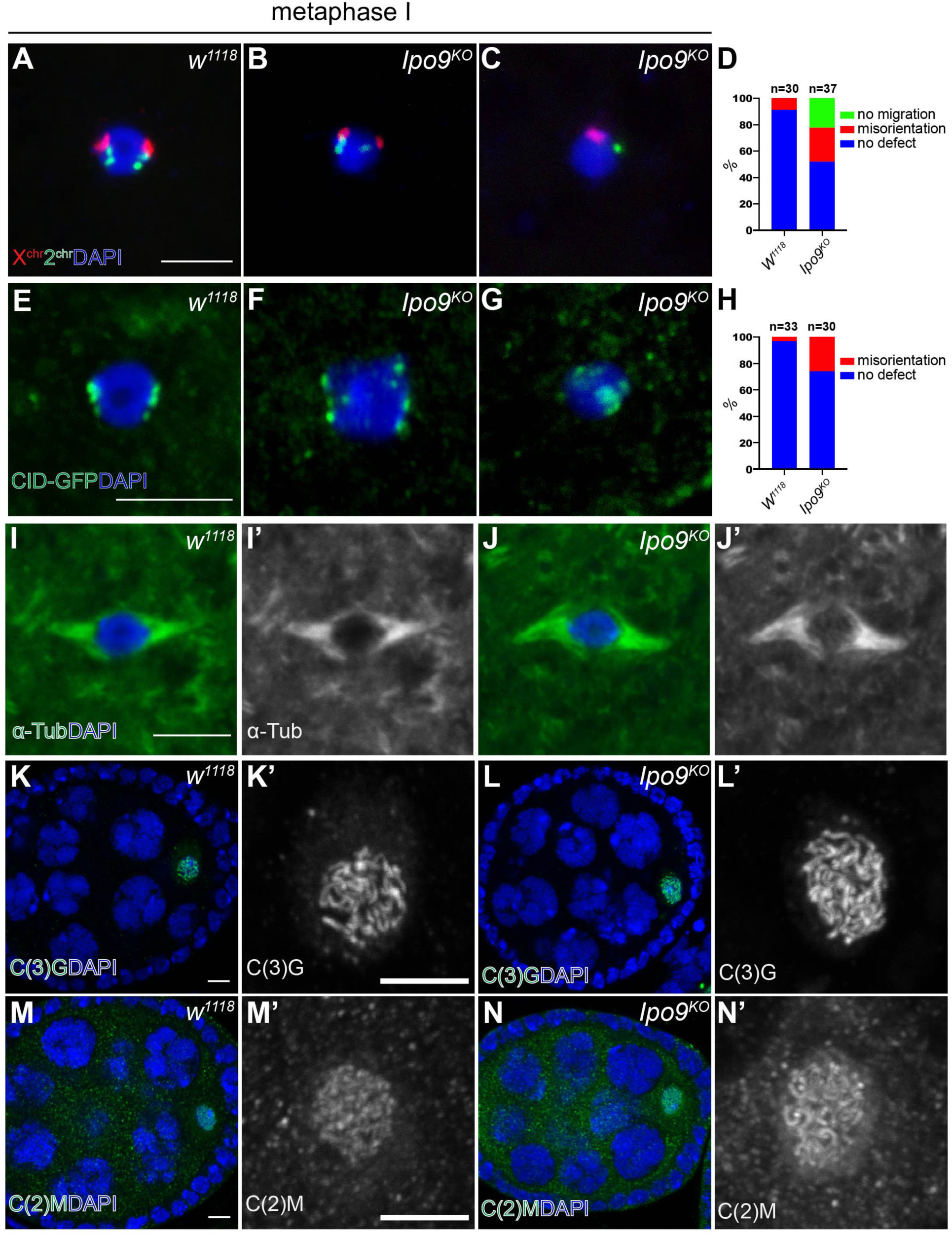
*Ipo9^KO^* oocytes at metaphase I show defects in chromosome orientation. (A-C) FISH using a X chromosome probe (red) and second chromosome probe (green) on oocytes in metaphase I, DAPI (blue). (A) *w^1118^* and (B-C) *Ipo9^KO^* oocyte. (D & H) Quantification of percentage of oocytes showing chromosome orientation defects. (E-G) Stained for CID-GFP (green) on oocytes at metaphase I, DAPI (blue) (E) control and (F-G) *Ipo9^KO^*oocyte. (I-J) Oocytes at metaphase I stained for α-Tub (green) and DAPI (blue). (I-I’) *w^1118^* and (J-J’) *Ipo9^KO^*. (K-L) Oocytes stained for C(3)G (green) at Stage 4 during oogenesis. (K-K’) *w^1118^* and (L-L’) *Ipo9^KO^*. (M-N) Oocytes stained for C(2)M (green). (M-M’) *w^1118^* or (N-N’) *Ipo9^KO^*. Scale bars 5μm.

Given the chromosome segregation defects we observed in *Ipo9* mutant female germ cells, we examined whether the nuclear import of meiotic specific machinery involved in sister chromosome pairing and DNA condensation was disrupted in the absence of *Ipo9*. Staining for the SC proteins, C(3)G and C(2)M, did not reveal any obvious differences between control and *Ipo9^KO^* ovarioles (Figure 3K-N’). As observed previously, we found that loss of *Ipo9* resulted in defects in nuclear actin accumulation (Figure S2) (Belin et al., 2015; Dopie et al., 2012; Kelpsch et al., 2016; Sokolova et al., 2018; Wineland et al., 2018). Determining the extent to which decreased levels of nuclear actin directly affect chromosome segregation or other aspects of meiosis represents important work for the future.

### Loss of Ipo9 causes male sterility

Similar to the phenotypes observed in females, no progeny were produced from matings between control females and *Ipo9* mutant males (Figure 4A). During spermiogenesis germ cell nuclei undergo dramatic shape changes to form needle-like structures. Close examination revealed that loss of *Ipo9* resulted in a failure of spermatid nuclei to change shape during spermiogenesis (Figure 4B-E). The clustered post-meiotic mutant nuclei remained round well beyond the stage during which they should have initiated changes in nuclear shape changes, resulting in the absence of mature sperm. We also compared sperm tail elongation and sperm individualization between control and *Ipo9^KO^* testes. Staining for α-Tubulin (α-Tub) to label the sperm tails does not reveal obvious differences between *w^1118^* and *Ipo9^KO^* testes (Figure S3A-B’). However, staining control and mutant testes using fluorescently labeled phalloidin (Cagan, 2003; Fabian and Brill, 2012), revealed that *Ipo9^KO^* testes do not form actin cones or waste bags (Figure S3C-F’), indicating that *Ipo9^KO^* spermatids do not go through individualization.

**Fig 4.**
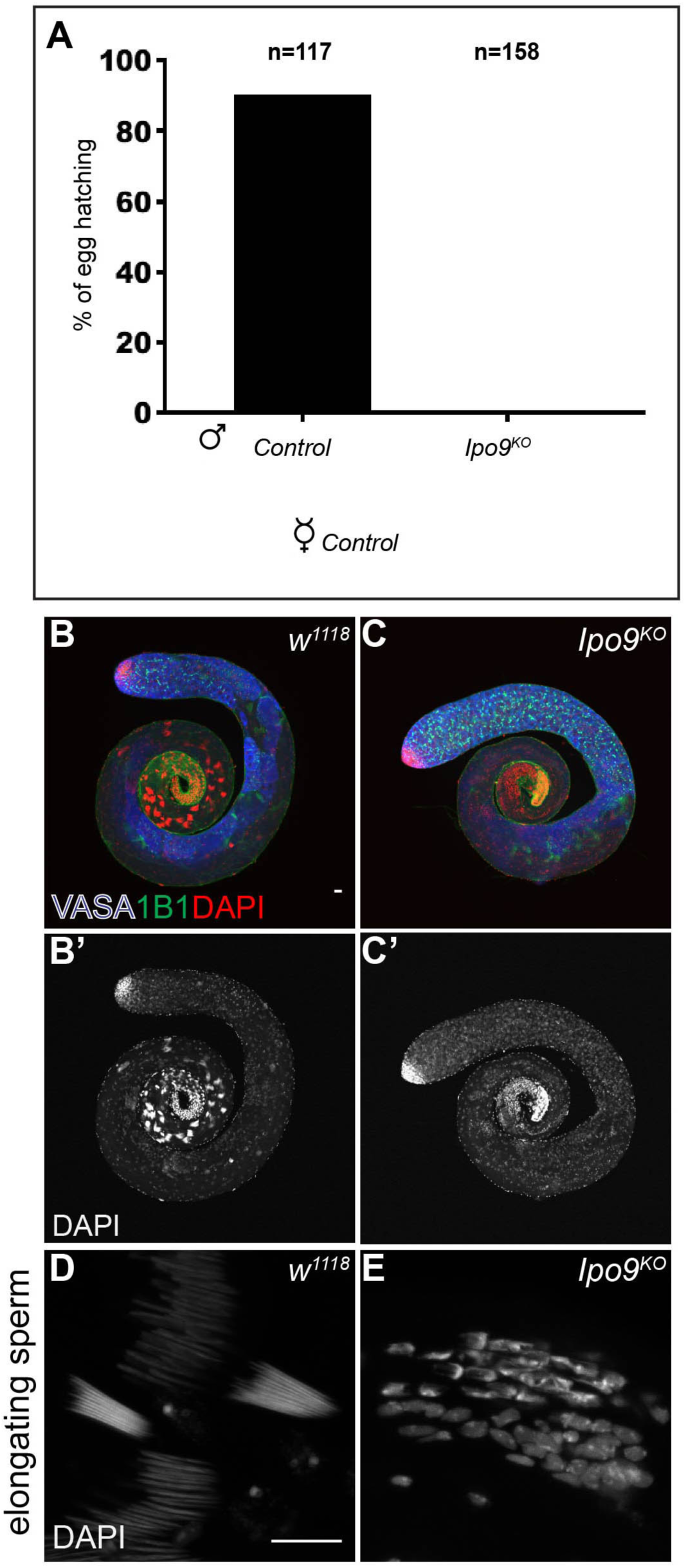
*Ipo9^KO^* males are unable to produce mature sperm. (A) Percentage of eggs that hatch after 5 days of being laid by *w^1118^* females crosses to *w^1118^* (control) or *Ipo9^KO^* males. (B-C’) *Drosophila* testes stained for VASA (blue), 1B1 (green) and DAPI (red). (B-B’) *w^1118^* (control) and (C-C’) *Ipo9^KO^*. (D,E) Cluster of elongating spermatids stained with DAPI. (D) *w^1118^* and (E) *Ipo9^KO^* testes. Scale bars 20μm.

We used both RNAi knockdown and cDNA rescue as independent methods to test whether the male phenotypes were caused specifically by loss of *Ipo9*. Driving *Ipo9* specific RNAi using germ cell specific drivers resulted in the same phenotypes in the testis as the *Ipo9^KO^* mutant (Figure S3G-I), supporting the idea that Ipo9 functions during male gametogenesis. Driving the full-length *Ipo9* cDNA transgene in an *Ipo9* mutant background using *vasa-gal4* rescued many of the morphological defects we observed during spermatogenesis, including sperm head elongation, but the *Ipo9^ΔN^* transgene did not (Figure S3J-M). However, expression of the *Ipo9^FL^* transgene did not fully rescue the male sterile phenotype, and most of the maturing sperm continued to be immotile. Given the similarities between the *Ipo9^KO^* and RNAi induced phenotypes, we expect that the inability of the full-length transgene to fully rescue the male sterile phenotype is caused by the failure of the *vasa-gal4* driven *Ipo9* expression to completely recapitulate the late-stage endogenous expression pattern of the protein during spermatogenesis.

### Ipo9 functions during male meiosis

To begin characterize whether male germ cells exhibit meiotic defects similar to what we observe in females, we crossed a GFP-tagged mitochondrial marker into the *Ipo9* mutant background so that we could examine the morphology of the nebenkern immediately after the completion of meiosis II (White-Cooper, 2004). Co-labeling for the mitochondrial marker and DNA showed that *Ipo9* mutants often exhibited defects at the onion stage, marked by the appearance of variably sized nuclei and nebenkern (Figure 5A-B”). In addition, the chromatin of *Ipo9* mutant nuclei appeared less condensed than control nuclei at the same stage of development (Figure 5A”-B”).

**Fig 5.**
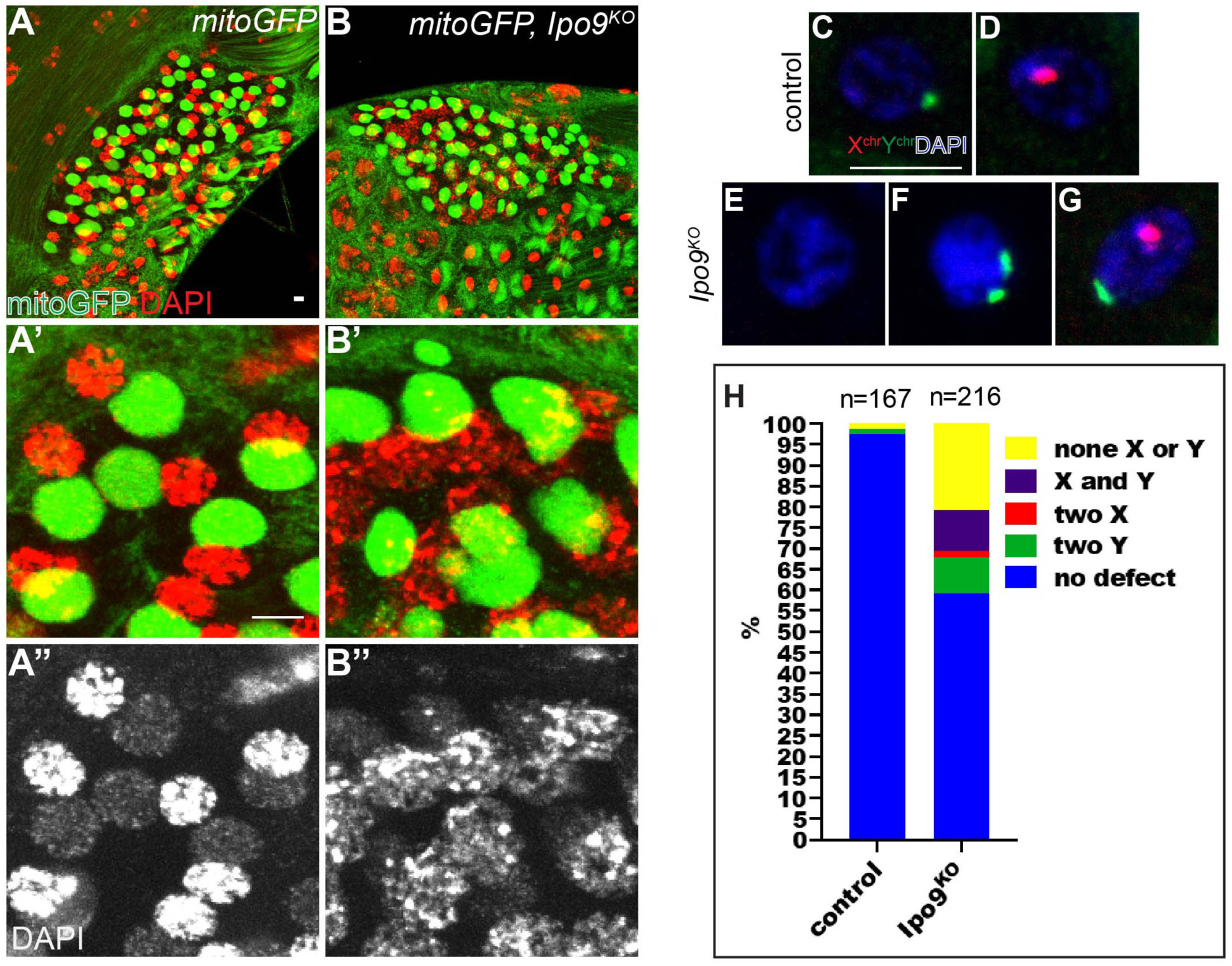
*Ipo9^KO^* spermatids exhibit chromosome segregation defects. (A-B) Spermatids at the onion stage stained for GFP (green) and DAPI (red). (A-A’’) control shows 1:1 ratio of condensed nuclei and rounded nebenkern. (B-B’’) *Ipo9^KO^* mutants exhibit nebenkern number and size defects. *Ipo9* mutant germ cells also display DNA condensation defects. (C-G) FISH using a X chromosome probe (red) and Y chromosome probe (green) and DAPI (blue) on spermatids at the onion stage. (C-D) control and (E-G) *Ipo9^KO^* spermatids. (H) Quantification of the percentage of spermatids showing chromosome segregation defects. Scale bars 5μm.

Next, we performed FISH experiments on wild-type and *Ipo9* mutant testes using probes specific for the X and Y chromosomes, focusing on the onion stage, just after the completion of meiosis II. As expected, half of the round spermatids in control samples were labeled with the probe for the X chromosome, while the other half carried a Y chromosome. By contrast, chromosome segregation defects were apparent in *Ipo9* mutant meiotic nuclei. We observed that 40% of *Ipo9* mutant spermatids contain neither a X nor a Y chromosome, both the X and Y chromosomes, two X chromosomes or two Y chromosomes at a stage when meiosis II should have been completed (Figure 5C-H). These results indicate that loss of *Ipo9* disrupts normal meiosis, in at least some fraction of male germ cells.

### Loss of *Ipo9* disrupts histone to protamine exchange in testes

As noted in our initial phenotypic characterization, *Ipo9* mutant spermatid nuclei remained round and failed to undergo the normal morphological changes that occur during the process of nuclear shaping. Shape changes in developing sperm occur as histones are being exchanged for protamines, but whether direct links between these processes exist remains unclear (Fabian and Brill, 2012). We examined whether histones were removed properly and replaced by protamines during the final stages of sperm development. Control spermatids showed replacement of the histone H2A and H2Av at the late elongation stage by protamine-B and overlapping of histone H2A or H2Av with protamine-B almost was never observed (Figures 6A, C-C’’’; S4A). By contrast, *Ipo9^KO^* spermatids accumulated nuclear protamine-B, in the presence of histone H2A and H2Av, neither of which was completely removed from germ cell nuclei (Figure 6 B, D-D’’’; S4B). Based on these results, we conclude that *Ipo9^KO^* spermatids have a defect in the chromatin packaging switch that marks mature sperm.

**Fig 6.**
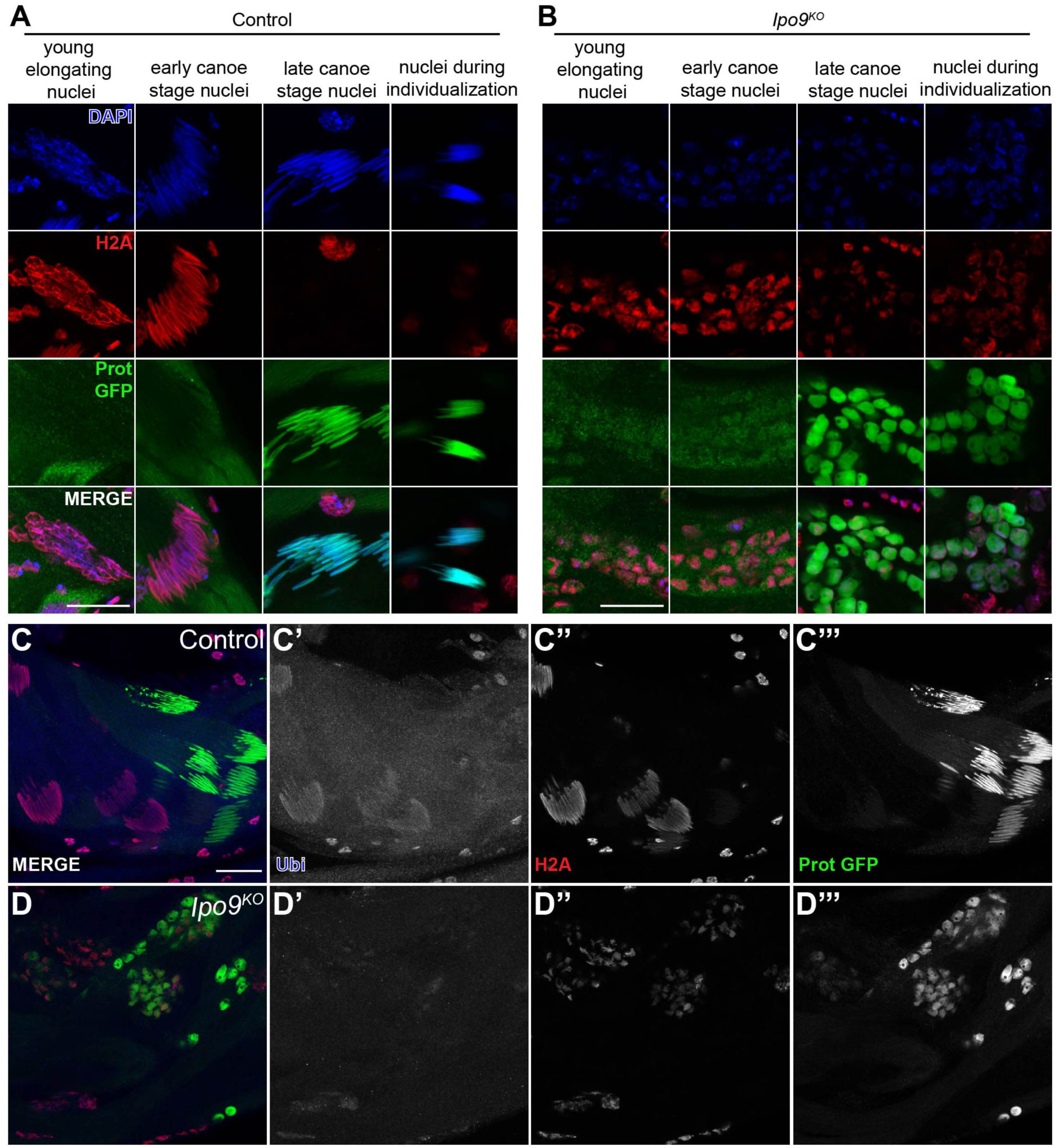
*Ipo9^KO^* spermatids show defect in H2A removal and histone ubiquitination. (A-B) Elongating nuclei stained for H2A (red), ProtB-GFP (green) and DAPI (blue). (A) *w^1118^* nuclei are able to elongate and replace histone with protamineB (B) *Ipo9^KO^* nuclei are unable to elongate and properly remove histones. (C-D) Testes stained for ubiquitin (blue), H2A (red) and protB-GFP (green). (C-C’’’) *w^1118^* control nuclei are positive for ubiquitination prior to protamine incorporation. (D-D’’’) *Ipo9^KO^* mutant nuclei show a reduction in ubiquitination. Scale bars 20μm.

The ubiquitin proteasome pathway has been implicated in histone degradation during spermiogenesis (Zhong and Belote, 2007). Because *Ipo9^KO^* spermatids have a defect in histone removal, we decided to explore whether histone ubiquitination is impaired in *Ipo9^KO^* testes. Staining for polyubiquitination in control testes showed spermatids positive for ubiquitination (Figure 6C-C’’’). However, nuclei that were in transition to protamine incorporation or had already accumulated protamines, were negative for polyubiquitination. Similar to control testes, *Ipo9^KO^* testes have germ cells that were positive for ubiquitination during early stages of sperm development (Figure 6D-D’’’). However, *Ipo9^KO^* spermatids were negative for ubiquitination at later stages, corresponding to when histones normally become ubiquitinated. Thus, loss of *Ipo9* appears to disrupt several of the changes to chromatin that occur during late sperm development.

### Ipo9 promotes the nuclear import of proteasome components during the late stages of sperm development

In an attempt to identify potential Ipo9 cargoes for nuclear import during male germ cell development, we immunoprecipitated Ipo9 from testes using the HA-tagged rescuing transgene under control of a *vasa-gal4* driver. Mass-spectrometry analysis revealed proteins that showed enrichment in the Ipo9 immunoprecipitation (IP) pellet versus the control IP pellet (Table S1). As noted above, ubiquitination plays a central role in removing histones from chromatin during the histone to protamine exchange that occurs during spermiogenesis (Rathke et al., 2007). However, the ubiquitin ligase responsible for this activity remains unknown. Interestingly, Ipo9 appears to associate with a number of ubiquitin ligases, including Hyperplastic Discs, CG5382, KLHL10, Sinah and CG31642 (Table S1). A couple of these gene exhibit sterility when mutated (Arama et al., 2007; Kaplan et al., 2010; Mansfield et al., 1994).

We also noted that several components of the proteasome, including Rpn1 and Rpt1 among others, appeared to associate with Ipo9. Previous efforts to define the global interactome of *Drosophila* proteins had also noted these same physical interactions (Guruharsha et al., 2011). To determine the functional significance of these results, we examined the sub-cellular distribution of several proteasome proteins, for which the necessary tagged transgenes have been developed, during the late stages of sperm development. This analysis showed that loss of *Ipo9* disrupts the normal nuclear import of Prosα3T, Prosα6T and Prosα2. For Prosα2, these defects could be observed immediately after meiosis II, and for all three were clearly evident during the canoe stage (Figures 7; S5,6), when nuclear shape changes occur and as histones are being replaced by protamines. These results indicate that Ipo9 plays a specific role in importing a number of nuclear factors that help to coordinate the chromatin re-organization that occurs late in *Drosophila* sperm development.

**Fig 7.**
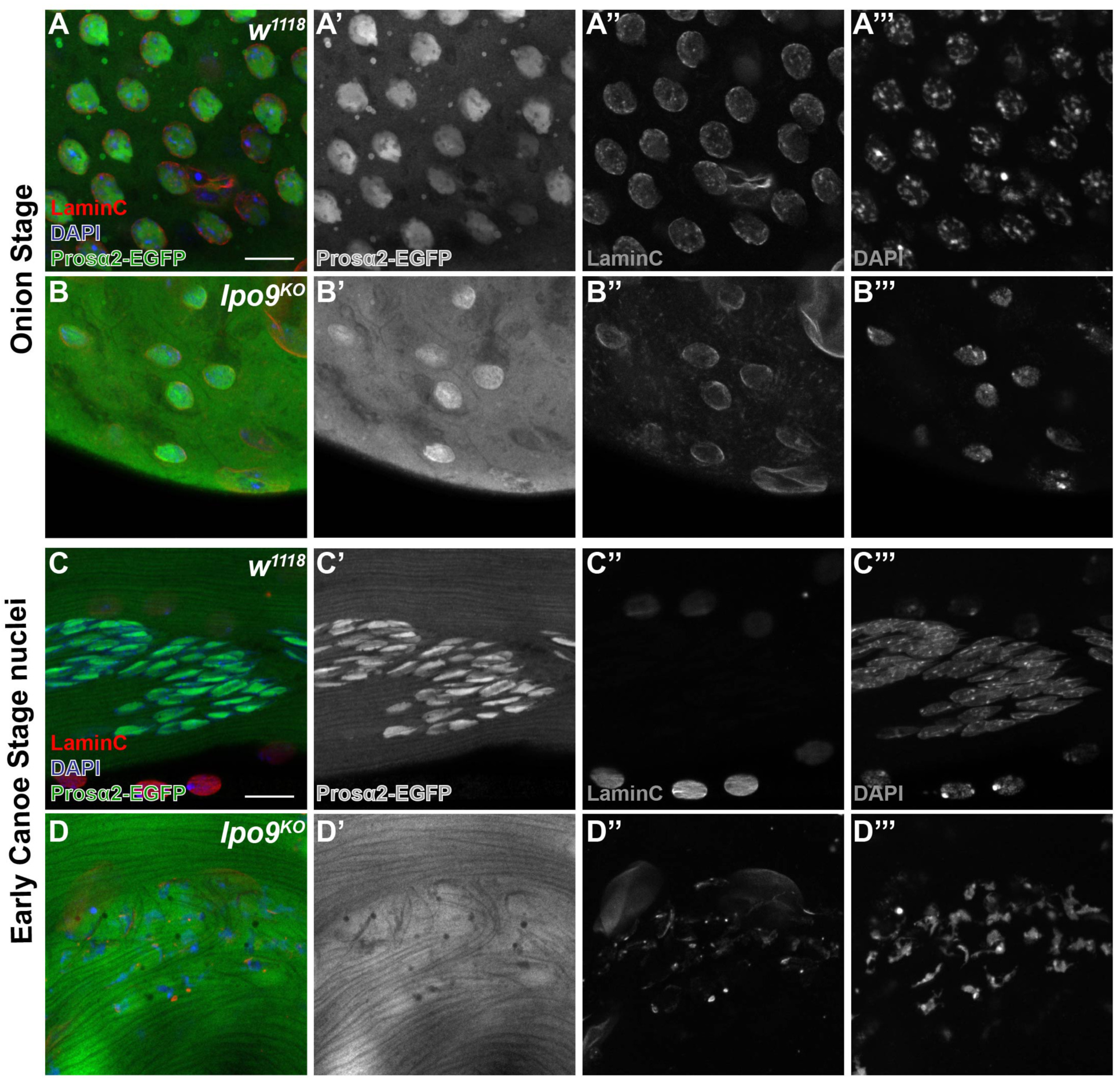
*Ipo9^KO^* spermatids show reduction of Prosα2 in the nucleus. (A-D) Spermatids at the onion stage and early canoe stage, stained for Prosα2-EGFP (green), LaminC (red) and DNA (blue). (A & C) *w^1118^* spermatids and (B & D) *Ipo9^KO^* spermatids. Scale bars 10μm.

## Discussion

Here, we provide evidence that *Ipo9* specifically regulates a number of critical processes during *Drosophila* gametogenesis. *Ipo9* null mutants survive to adulthood but exhibit female and male sterility. In the ovary, loss of *Ipo9* results in defects in chromosome orientation and segregation during meiosis, resulting in mitotic catastrophes during early embryogenesis in progeny derived from *Ipo9* homozygous mutant females. *Ipo9* mutant males also exhibit numerous phenotypes during germ cell development including defects in meiosis, and disruption of the nuclear shape changes and failure to fully exchange histones for protamines during spermiogenesis. Together, these represent a unique spectrum of phenotypes when compared to other *Drosophila* β-karyopherin family members. Of the 12 *Drosophila* β-karyopherin genes that have been genetically characterized, loss-of-function alleles in 7 result in lethality (Baker et al., 2002; Collier et al., 2000; Giagtzoglou et al., 2009; Higashi-Kovtun et al., 2010; Ilius et al., 2007; Jackel et al., 2015; Jakel and Gorlich, 1998; Kahsai et al., 2016; Lippai et al., 2000; Natalizio and Matera, 2013; Tekotte et al., 2002; VanKuren and Long, 2018). Several other importin mutants do survive until adulthood, including *ebo^mut^*, *apl^null^* and *arts^null^*. *ebo^mut^* homozygotes display neuronal defects, *apl^null^* mutants are male sterile, while *arts^null^* mutant females produce smaller eggs that cannot be fertilized (Collier et al., 2000; Ilius et al., 2007; VanKuren and Long, 2018). In addition, a *ketel* dominant negative mutant (*ketel^D^*) shows female sterile phenotype and embryos derived from these flies exhibit chromosome segregation defects somewhat similar to those displayed by *Ipo9^KO^* mutants (Schupbach and Wieschaus, 1991; Timinszky et al., 2002; Tirian et al., 2000). Thus, amongst *Drosophila* karyopherin family members studied to date, *Ipo9* is the only gene that displays specific defects during meiosis in both females and males, and in late sperm development when mutated.

Transgenic rescue experiments confirm that Ipo9 functions to promote the transport of molecules from the cytoplasm to the nucleus during oogenesis and spermatogenesis. A full-length Ipo9 transgene rescues most of the sterile phenotypes exhibited by *Ipo9* mutants when driven in the germline. We suspect the failure of the Ipo9 wild-type transgene to fully rescue the male sterility of the mutant is likely due to the failure of the *vasa-gal4* driver to fully recapitulate the endogenous expression pattern of *Ipo9*. The N-terminal domains of β-karyopherin proteins normally promote cytoplasmic-to-nuclear trafficking by contacting the nuclear pore and helping cargoes move through the nuclear pore complex. This domain also binds to RanGTP, and thus participates in the cycling of importins back-and-forth between the cytoplasm and nucleus (Bange et al., 2013; Chi and Adam, 1997; Fried and Kutay, 2003; Kutay et al., 1997; Strom and Weis, 2001). Strikingly, deletion of the N-terminal karyopherin domain renders the transgene non-functional, confirming that Ipo9 acts as an essential transport factor during gametogenesis in both males and females.

The transition from histone-based to protamine-based chromatin organization is essential for the nuclear shaping that leads to a highly compact sperm nucleus (Rathke et al., 2014). *Ipo9^KO^* nuclei are able to incorporate protamine-B, however histone H2A and H2Av are not completely removed. These results may partially explain why *Ipo9^KO^* nuclei do not elongate properly. Evidence of histone ubiquitination prior to transition to protamine-based incorporation and delay in histone removal in a proteasome component mutant suggest that the ubiquitin proteasome pathway is involved in histone removal in spermiogenesis (Awe and Renkawitz-Pohl, 2010; Zhong and Belote, 2007). Interestingly, *Ipo9^KO^* nuclei do not exhibit strong nuclear ubiquitination after protamine incorporation, even though they still retain nuclear histones (Figure 6D-D’’’). Additionally, we observed that *Ipo9^KO^* spermatids showed a significant reduction in the nuclear localization of several proteasome proteins, including Prosα6T, Prosα3T and Prosα2, compared to the control spermatids. These results suggest that the ligase(s) responsible for histone ubiquitination and components of proteasome that ultimately degrades ubiquitinated histones are both potential cargoes of Ipo9. Interestingly, Ipo9 appears to physically associate with a number of specific ubiquitin ligases, three of which have been implicated in the regulation of male germ cell development (Arama et al., 2007; Kaplan et al., 2010; Mansfield et al., 1994), and with several components of the proteasome. Perhaps Ipo9 has evolved to the temporally coordinate the import these functionally related proteins during late sperm development. Such specialization in nuclear import may offer an economy of scale that wouldn’t exist if the responsibility of nuclear import during this critical phase of sperm development, when the cytoplasm and nuclei of sperm are becoming highly compacted, were spread across a number of potentially redundant β-karyopherins. This type of coordination in trafficking has been proposed previously in different contexts (Bange et al., 2013). Thus, the further study of Ipo9 cargoes during sperm development may reveal critical unknown factors that play roles in meiosis, chromosome compaction and segregation, and nuclear shape changes.

## Acknowledgements

We would like to thank K. McKim, Y. Yamashita, J. Belote, R. Glaser, the Bloomington *Drosophila* Stock Center and the Developmental Studies Hybridoma Bank for reagents. We thank previous and current members of the Buszczak lab for comments and advice. This work has been funded by NIH/NIGMS through R01GM116885 to T.L.T. and R01GM125812 to M.B.

## Author Contributions

V.P. and G.C.K. conducted the experiments. T.L.T and M.B. designed experiments. V.P. and M.B. wrote the manuscript. V.P. T.L.T. and M.B edited the manuscript.

## Material and Methods Fly Stocks

Fly stocks were maintained at 22^0^C–25^0^C on standard cornmeal-agar-yeast food unless otherwise noted. RNAi knockdown in male flies was achieved at 29^0^C. The following stocks were used in this study: *w^1118^* (BL-6326), *His2Av-mRFP1* (BL-34498), *ProtamineB-eGFP* (BL-58406), *Mat-*α*-Tub-gal4* (BL-80361 II^chr^ and III^chr^), *MTD-gal4* (BL-31777), UAS- *Ipo9^RNAi^* (BL-33004), *sqh-EYFP-Mito* (BL-7194). *UASp-HA-Ipo9^FL^* and *UASp-HA-Ipo9^ΔN^* was inserted into attP40(BL-25709) using phiC31 integrase (Rainbow Transgenics). *vasa-gal4* was a gift from Y. Yamashita. *Prosα6T-EGFP*, *Prosα3T-EGFP* and *Prosα2-EGFP* were gifts from Dr. John Belote.

### Cloning *Ipo9*

RNA was extracted from *w^1118^* ovaries and made into cDNA using a SuperScript II-Strand Kit (Life Technologies). We next performed PCR using Ipo9^FL^ specific primers (F5’CACCATGTCGCTGCAATTCCAAAACG and R5’CTACTTCTGCTGGACCTTGCTG) To generate Ipo9^ΔN(36-144aa)^ we performed PCR using these primers (F5’ CACCATGTCGCTGCAATTCCAAAACG and R5’TTCTGTCTGCTGCAGGACTCC first & R5’GAGGAGCGTATCTTTGAATTGGGTTCTGTCTGCTGCAGGACTCC second) for fragment 1 and (F5’CCCAATTCAAAGATACGCTCCTC and R5’CTACTTCTGCTGGACCTTGCTG) to generate fragment 2. Then PCR SOE was performed to stitch fragment 1 with fragment 2. PCR products were cloned into pENTR (Life Technologies) and swapped into pAHW (Drosophila Gateway Vector Collection) using an LR reaction.

### Generating the *Ipo9^KO^* allele

To generate the *Ipo9^KO^* allele, guide RNAs were designed using http://tools.flycrispr.molbio.wisc.edu/targetFinder (Guide1 5’CTTCGCGCTATCACATGTAGTCAA/5’AAACTTGACTACATGTGATAGCGC and Guide2 5’CTTCGGTGGACAGAAAGTTGAGTA/5’AAACTACTCAACTTTCTGTCCACC) and synthesized by IDT as 5’ unphosphorylated oligonucleotides, annealed, phosphorylated, and ligated into the BbsI sites of the pU6-BbsI-chiRNA plasmid (Gratz et al., 2013). Homology arms were PCR amplified and cloned into pHD-dsRed-attP (Gratz et al.,2014) (arm1F5’GCTACACCTGCATGCTCGCGTTCATGTGCAAGCGCAAGTC, R5’GTCACACCTGCACTGCTACAACGGGCGTTTTGCAAGACTG arm2 F5’CGTAGCTCTTCGTATCAACTTTCTGTCCACCGTTCC, R5’CGATGCTCTTCCGACGCGAACCGAATCGTAACTGGC)(Addgene). The pHD-dsRed-attP vector was cut with the enzymes AarI and SapI. Guide RNAs and the donor vector were co-injected into nosP Cas9 attP40 embryos at the following concentrations: 250 ng/ml pHDdsRed-attP donor vector and 20 ng/ml of each of the pU6-BbsI-chiRNA plasmids containing the guide RNAs (Rainbow Transgenics).

### PCR verification of *Ipo9^KO^*

Primers for PCR1 Ipo9Aar1outF5’ CAAGCCGCAAATGATGCTGCTG and DsRedstartR5’ CATGAACTCCTTGATGACGTCCTC. PCR2 DsRedendF5’GACTACACCATCGTGGAGCAG and Ipo9Sap1outR5’CTTTGCCTTTGGCTCAGAGAAGC. Internal primers Exon2F5’GGAACTGGGTCCAGTAGTCATAC3’ and Exon5R5’GAGGTGGAGATTCTTGATGCAC3’.

### Immunofluorescent staining in ovaries, testes and embryos

Ovaries and testes were dissected in Grace’s Medium. Ovaries and testes were fixed for 10 minutes with gentle rocking in 4% formaldehyde in PBS. Fixed ovaries and testes were briefly rinsed three times and permeabilized in PBST (1X PBS + 0.3% Triton X-100) at room temperature for 1hr before adding primary antibody.

*Drosophila* embryos were stained according to (Mani et al., 2014). Embryos were dechorionated in 50% bleach for 2-3mins. Then embryos were rinsed in 1X PBS 2 times. Embryos were fixed in 50% heptane and 50% fixative solution (3 parts fixative solution, 1.33X PBS and 67 mM EGTA:1part 37% formaldehyde) for 10min. After fixation, the aqueous phase (bottom) was removed and replaced with an equal volume of 100% methanol. Then the embryos were vortexed rigorously for 1-2mins. Embryos were rinsed with 100% methanol 2 times. Then embryos were either stored at −20°C or rehydrated. To rehydrate, embryos were washed in a series of 70%MeOH: 30%PBST, 50%MeOH: 50%PBST, 30%MeOH:70% PBST and finally 100% PBST for 20 min each. Then embryos were blocked in 5% normal goat serum for 1hr at RT.

Incubation with primary antibody was in 3% bovine serum albumin (BSA) in PBST at 4 °C at least for 20hrs. Samples were washed three times for 20 min in PBST, incubated with secondary antibody in 3% BSA in PBST at room temperature for 3–5 hrs and then washed three times 20 min each in PBST. Samples were mounted in VectaShield mounting medium with DAPI (Vector Laboratories). The following antibodies were used (dilutions noted in parentheses): mouse anti-Hts (1B1) (1:20), rat anti-VASA(1:20), mouse actin-JLA20(1:10) and LaminC (LC28.26) (1:10) (Developmental Studies Hybridoma Bank, Iowa), rabbit anti-Vasa-d-260 (1:200 Santa Cruz), mouse actin-C4(1:100 MAB1501 Millipore Sigma) rat anti-HA 3F10 (1:100; Roche), rabbit anti-GFP(1:1000 Molecular Probes), rat α-Tub (1:100 YL1/2 Abcam), chicken anti-GFP (1:1000 Novus Biological) mouse anti-ubiquitin (1:100 P4D1 Cell Signalling), Rabbit anti-RFP (1:1000 Rockland) rabbit anti-H2A(1:2000, from Dr. Robert L. Glaser Lab), rabbit anti-C(3)G (1:1000, from Dr. Kim Mckim), rabbit anti-C(2)M (1:1000, from Dr. Kim Mckim), rhodamine phalloidin (1:200, R415 300U Invitrogen); Cy3, Cy5, FITC (Jackson Laboratories) or Alexa 488 (Molecular Probes) fluorescence-conjugated secondary antibodies were used at a 1:200 dilution. Images were taken using a Zeiss LSM800 confocal microscope with a 40× oil immersion objective and processed using Image J.

### Fertility Assays

3-7day old males and virgin females of the appropriate genotype were mated in mating cages with grape juice (3%) agar plates with a little bit of wet yeast. The flies were allowed to lay eggs for 12-24hrs at 22-25^0^C.

### Western blotting

For protein extraction, ovaries from fatten flies were dissected in Grace’s medium, physically disrupted and extracted with sample buffer with 20% BME using pestle followed by heating at 90°C for 10 minutes. Protein electrophoresis and wet transfer systems were used. After running the SDS-PAGE gel, the proteins were transfer to an Amersham Hybond ECL nitrocellulose membrane (GE Healthcare, RPN2020D). For blotting, the following primary antibodies were used in fresh PBST buffer (1XPBS with 0.1%Tween20 and 5% Biorad non-fat milk): mouse anti-ActinJLA20(1:100) and rat anti-VASA(1:10000) from Developmental Studies Hybridoma Bank, Iowa, mouse anti-ActinC4 (1:1000 MAB1501 Millipore Sigma) and mouse anti-HA (1:1000 5B1D10 ThermoFisher). After overnight incubation at 4^0^C, the membranes were washed for 20mins three times in PBST buffer without milk before incubating with secondary antibodies for 2 hours at RT. HRP-conjugated anti-mouse and anti-rat secondary antibodies (Jackson Laboratories) were used at a 1:2000 dilution. After incubation with the secondary antibody, the membranes was washed three times for 20 min each and then incubated with ECL Western Blotting Detection Reagents (GE Healthcare, RPN2106).

### Oocytes preparation for meiosis I

The following protocol was adopted from (Radford and McKim, 2016). Females fed of wet yeast were aged for three to five days at 25^0^C to enrich for oocytes in metaphase. Ovaries were dissected in 1X PBS solution at RT. For fixation, 687.5 µl of Fixation Buffer (1X PBS, 150 mM sucrose) was freshly mixed with 312.5 µl 16% formaldehyde. 0.5 ml of this solution was added to the ovaries and incubated for 2.5 mins on a nutator. 0.5 ml heptane was then added to top of the fix solution and vortexed for 1 min. The tissue was then allowed to settle for 1 min. The fixative was removed and 1 ml of 1X PBS was added to the sample and vortexed for 30 sec. Samples were allowed to settle for 1 min, before another quick wash with 1X PBS. To remove the membranes, 3 to 4 pairs of ovaries were added to a glass slide. The ovaries were then separated into individual ovarioles using forceps. 1X PBS was then added as necessary to prevent the ovarioles from drying out. A coverslip was placed on top of the ovarioles and gently “rolled” until all membranes were removed. The samples were then subjected to immunofluorescent staining or FISH.

### FISH

A protocol adapted from the Fox lab was used with oligopaints (Beliveau et al., 2014; Beliveau et al., 2015; Beliveau et al., 2012). Oocytes were prepared for examining meiosis I according to (Radford and McKim, 2016). Testes were dissected in Grace’s media and fixed in 4% paraformaldehyde buffered in 1X PBS for 10 minutes. The samples were then washed once in 1X PBS for 1 min, 1X in PBS + Tween (100 µl Tween for every 100 ml 1X PBS) for 1 min, 1X in PBS + Triton (250 µl Triton for every 50 ml 1X PBS) for 10 min, 1X in PBS + Tween (100 µl Tween for every 100 ml 1X PBS) for 1 min and finally in 0.1N HCL for 5 min. The samples were then washed three times in 2X SSCT for 2 min, once in 2X SSCT/50% Formamide for 5 min and once in 2X SSCT/50% Formamide at 60^0^C for 20 min. During the final wash, the oligopaint probe (200-300 pmol) was added hybridization mix (12.5 µl 2x hyb cocktail (10ml-4ml 50% dextran sulfate solution, 2ml 20X SSC, 4ml ddH2O), 12.5 µl formamide, 1 µl 10mg/ml RNase) and mixed by vortexing and then spun down. Protect from light until needed. This mixture was added to each sample and then incubated at 78^0^C for 2.5 minutes. The samples were then incubated in a 42^0^C water bath overnight. The samples were washed in 2X SSCT/50% formamide at 60^0^C for 1 minute. After this wash, the samples were moved to room temperature and washed three times in 2X SSCT/50% formamide for 10 min and three times in 0.2X SSC for 10 min. The samples were then mounted in Vectashield with DAPI.

The following fluorescently labeled oligos (IDT) were used for FISH: X-5’Cy3-TTTTCCAAATTTCGGTCATCAAATAATCAT Y-5’Alexa488-N/AATACAATACAATACATTACAATACAATAC 2-5’Cy5-AACACAACACAACACAACACAACACAACAC

### Immunoprecipitation followed by Mass Spectrometry

Preparation of crude protein lysate from fly adult testes. 250 pairs of testes from *Vasa-gal4>+* and *Vasagal4>UASp-HA-Ipo9^FL^* flies were dissected in cold PBS. Testes were washed twice in PBS before lysed on ice in lysis buffer (50mM Tris pH8.0, 137mM NaCl, 1mM EDTA, 1% Triton X-100, 10% glycerol, 10mM NaF and protease inhibitors). After centrifugation, the supernatants were incubated with rat anti-HA (Affinity Matrix Roche) 3-6hrs at 4°C. The beads were then quickly washed 3 times with lysis buffer and boiled in Laemmlli sample buffer with BME. Samples were submitted to the UT Southwestern proteomics core for complex mixture trypsin service.

---

**Fig S1 (Related to Figure 1).**
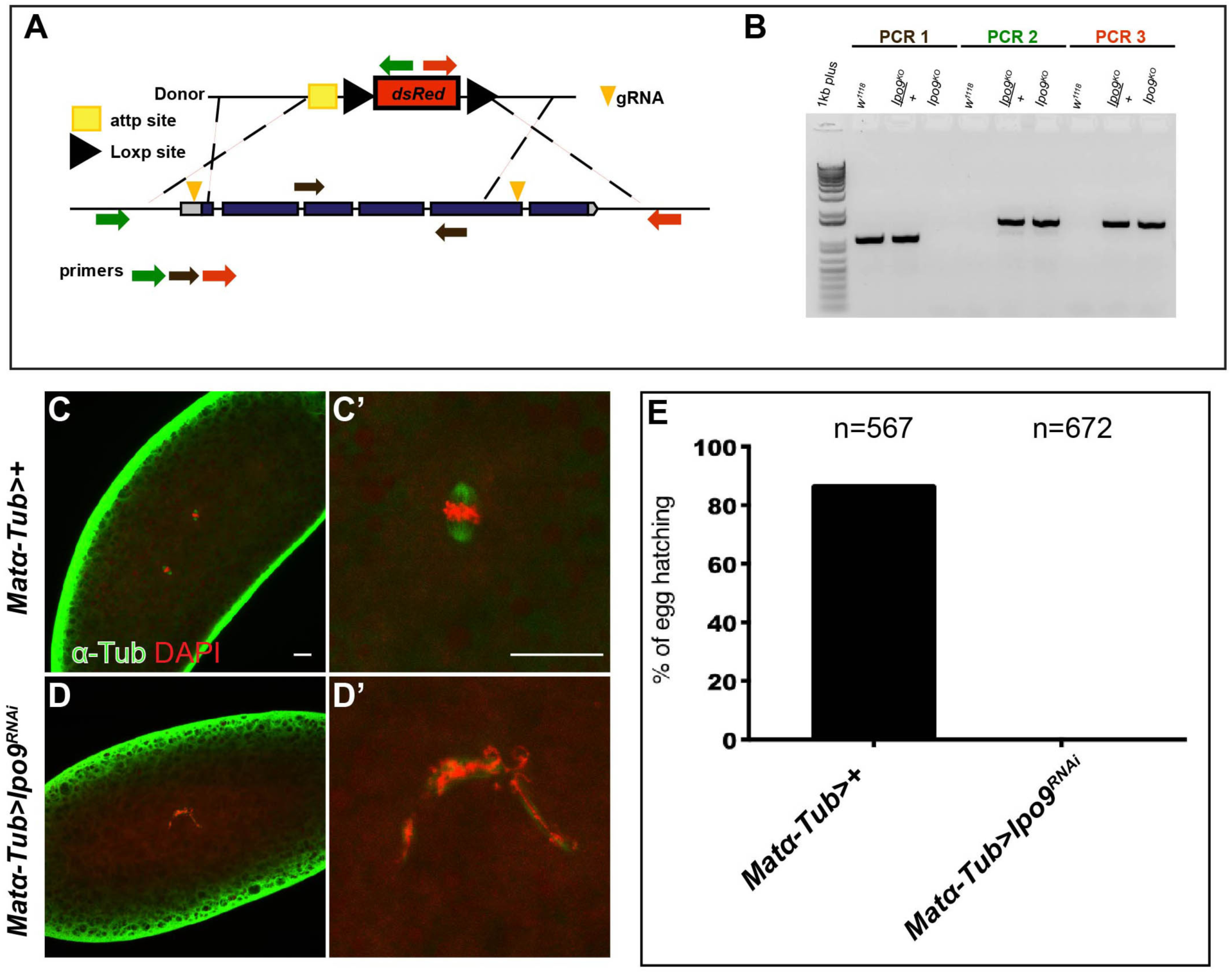
Loss of *Ipo9* in germ cells results in fertility defects. (A) Schematic of strategy used to knock-in 3xP3-dsRed cassette to replace the majority of *Ipo9* sequence. (B) PCR verification of knock-in of *3xP3-dsRed* cassette into the *Ipo9* locus in the *Ipo9^KO^* allele. (C-C’) Embryos from *Mat-*α*-Tub*-*gal4* (control) and (D-D’) *Mat-*α*-Tub*-*gal4>Ipo9^RNAi^* mothers stained for α-Tub (green) and DAPI (red). (E) Percentage of eggs that hatch after 5 days of being laid by *Mat-*α*-Tub*-*gal4* (control) and *Mat-*α*-Tub*-*gal4 >Ipo9^RNAi^* females crossed to *w^1118^* males. Scale bars 20μm.

**Fig S2 (Related to Figure 3).**
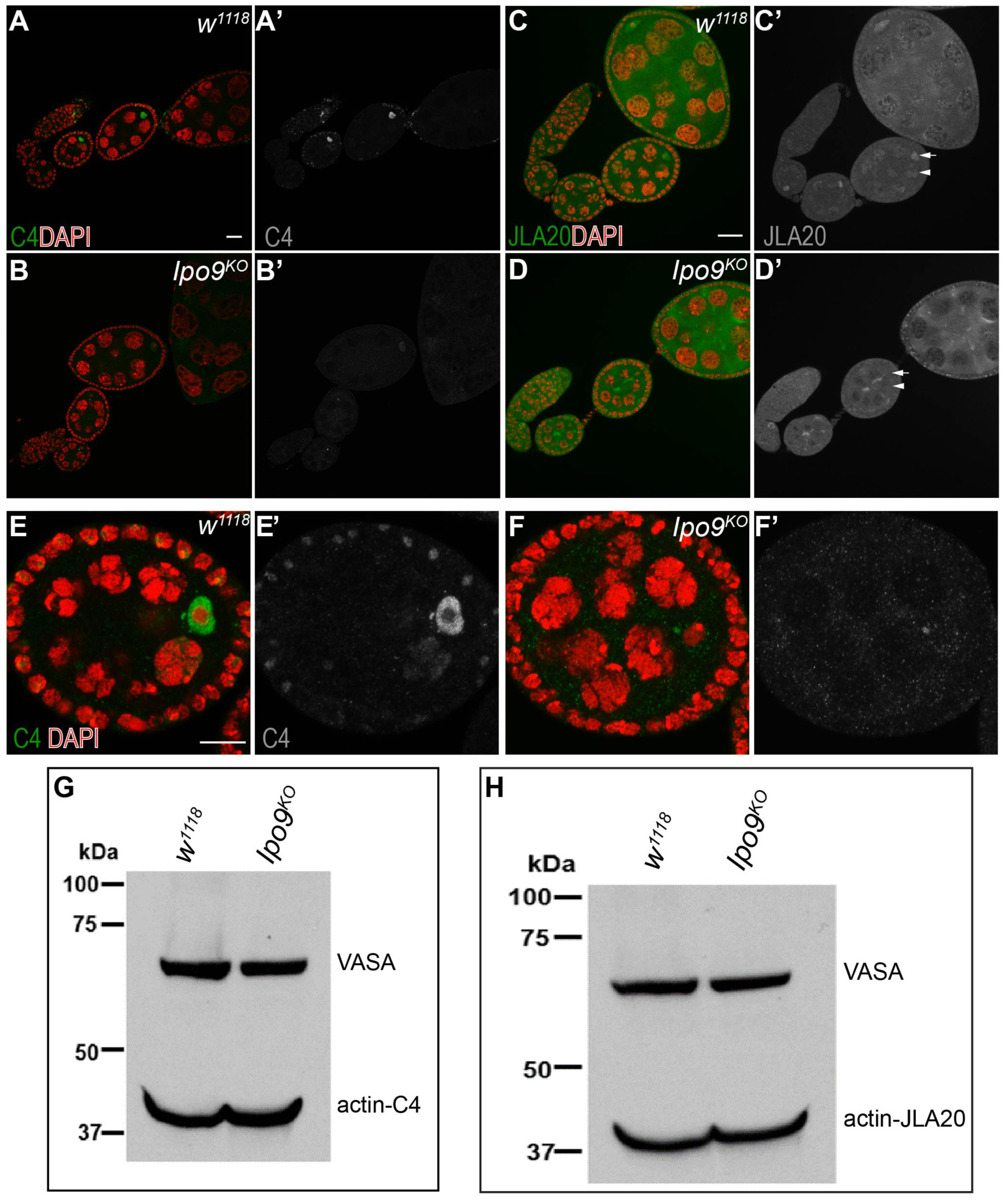
*Ipo9^KO^* germ cells show a reduction of nuclear actin. (A-B) Ovarioles stained for actinC4 (green) DAPI (red). (A-A’) *w^1118^* (control) and (B-B’) *Ipo9^KO^*. (C-D) Stained for ActinJLA20 (green) and DAPI (red). (C-C’) *w^1118^* (control) and (D-D’) *Ipo9^KO^* ovariole. Scale bars 20μm. (E-F) Egg chambers Stage 4 stained for actinC4 (green) and DAPI (red). (E-E’) *w^1118^* (control) and (F-F’) *Ipo9^KO^*. Scale bars 10μm. (G) Western blot showing Actin (C4) and Vasa expression from and *w^1118^* (control) and *Ipo9^KO^* ovaries. (H) Western blot from showing Actin (ActinJLA20) and Vasa expression from *w^1118^* (control) and *Ipo9^KO^* ovaries.

**Fig S3. (Related to Figure 4).**
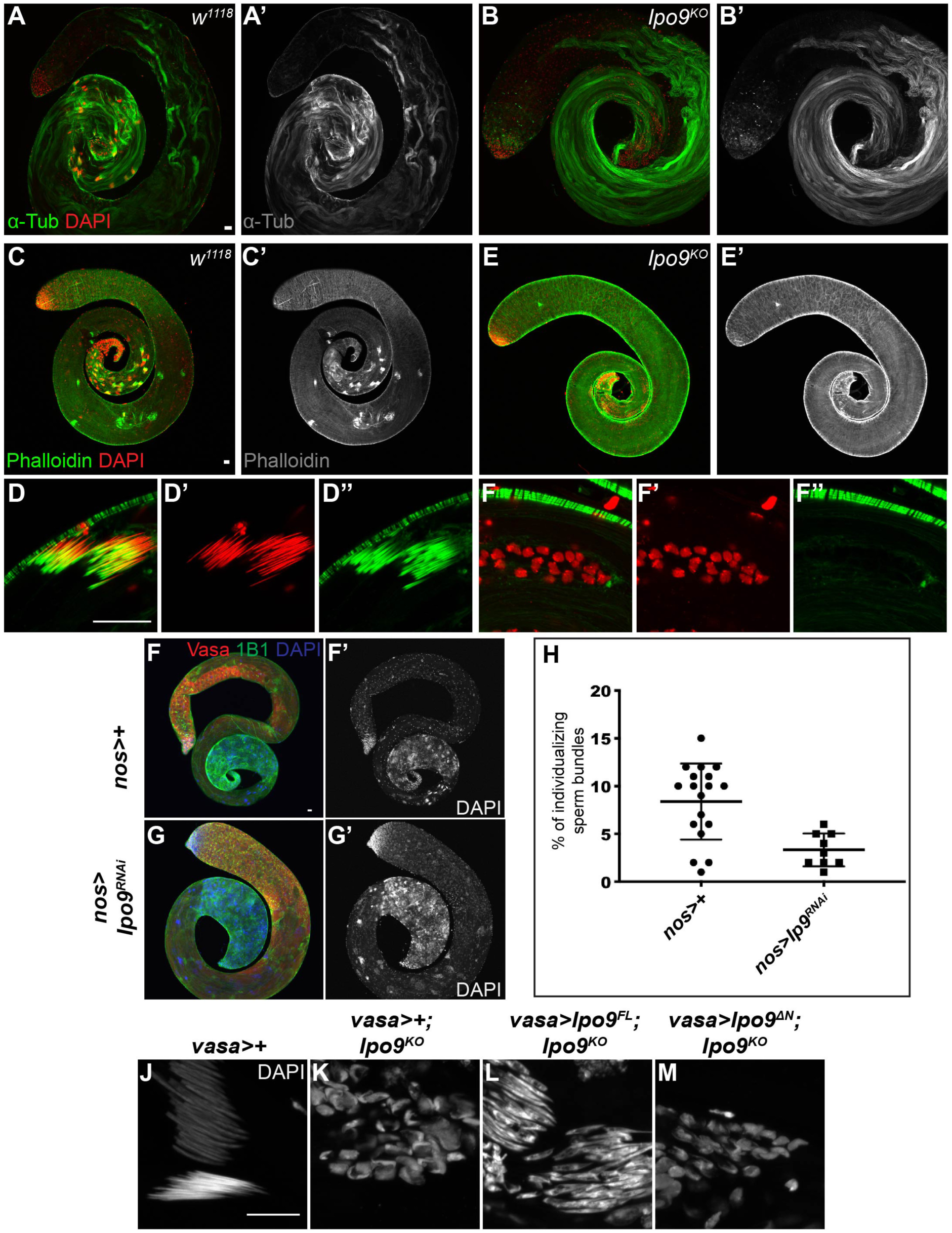
*Ipo9^KO^* nuclei are unable to individualize. (A-B’) Testes stained for α-Tub (green) and DAPI (red). (A-A’) *w^1118^* testis and (B-B’) *Ipo9^KO^* testis. (C-F”) Testes stained for phalloidin (green) and DAPI (red). (C-D) *w^1118^* testis and nuclei and (E-F’) *Ipo9^KO^* testis and nuclei. Scale bars 20μm. (G-G’) A *MTD-gal4* (control) and (H-H’) *MTD-gal4*>*Ipo9^RNAi^ Drosophila* testes stained for VASA (red), 1B1 (green) and DAPI (blue). (I) Number of individualizing sperm per testis from *MTD-gal4* (control) and *MTD-gal4*>*Ipo9^RNAi^* males. Scale bars 20μm. (J-M) Elongating sperm stained with DAPI (gray) for these genotypes (J) *vasa-gal4>;+*, (K) *vasa-gal4>;Ipo9^KO^*, (L) *vasa-gal4>Ipo9* ^FL^*;Ipo9^KO^* and (M) *vasa-gal4>Ipo9*^ΔN^*;Ipo9^KO^*. Scale bars represent 10µm.

**Fig S4 (Related to Figure 6).**
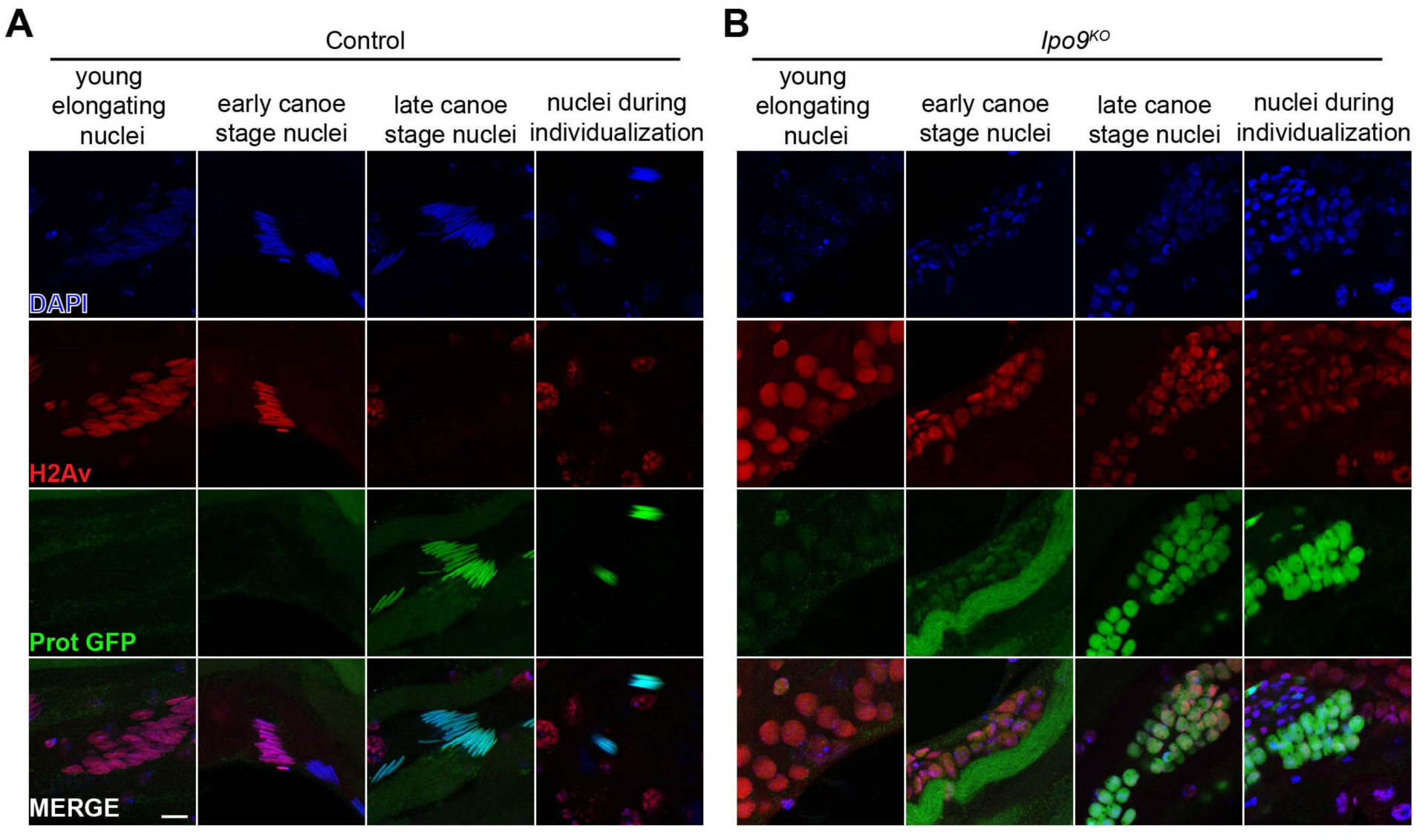
*Ipo9^KO^* spermatids show defect in H2Av removal. (A-B) Elongating nuclei stained for H2Av (red), ProtB-GFP (green) and DAPI (blue). (A) *w^1118^* nuclei are able to elongate and replace histone with protamineB. (B) *Ipo9^KO^* nuclei are unable to elongate and properly remove histones. Scale bars 10μm.

**Fig S5 (Related to Figure 7).**
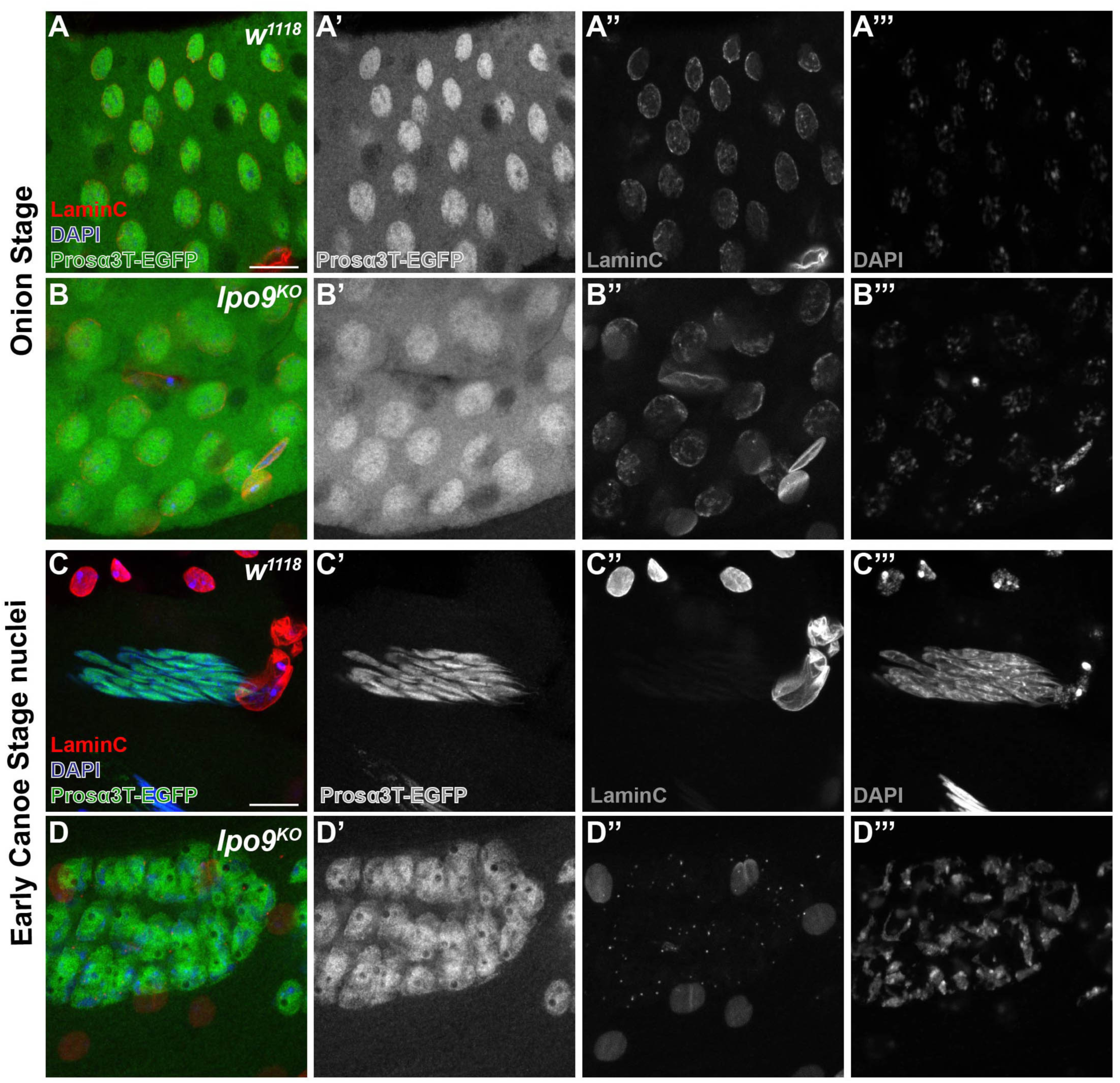
*Ipo9^KO^* spermatids show reduction of Prosα3T in the nucleus. (A-D) Spermatids at the onion stage and early canoe stage, stained for Prosα3T-EGFP (green), LaminC (red) and DNA (blue). (A & C) *w^1118^* spermatids and (B & D) *Ipo9^KO^* spermatids. Scale bars 10μm.

**Fig S6 (Related to Figure 7).**
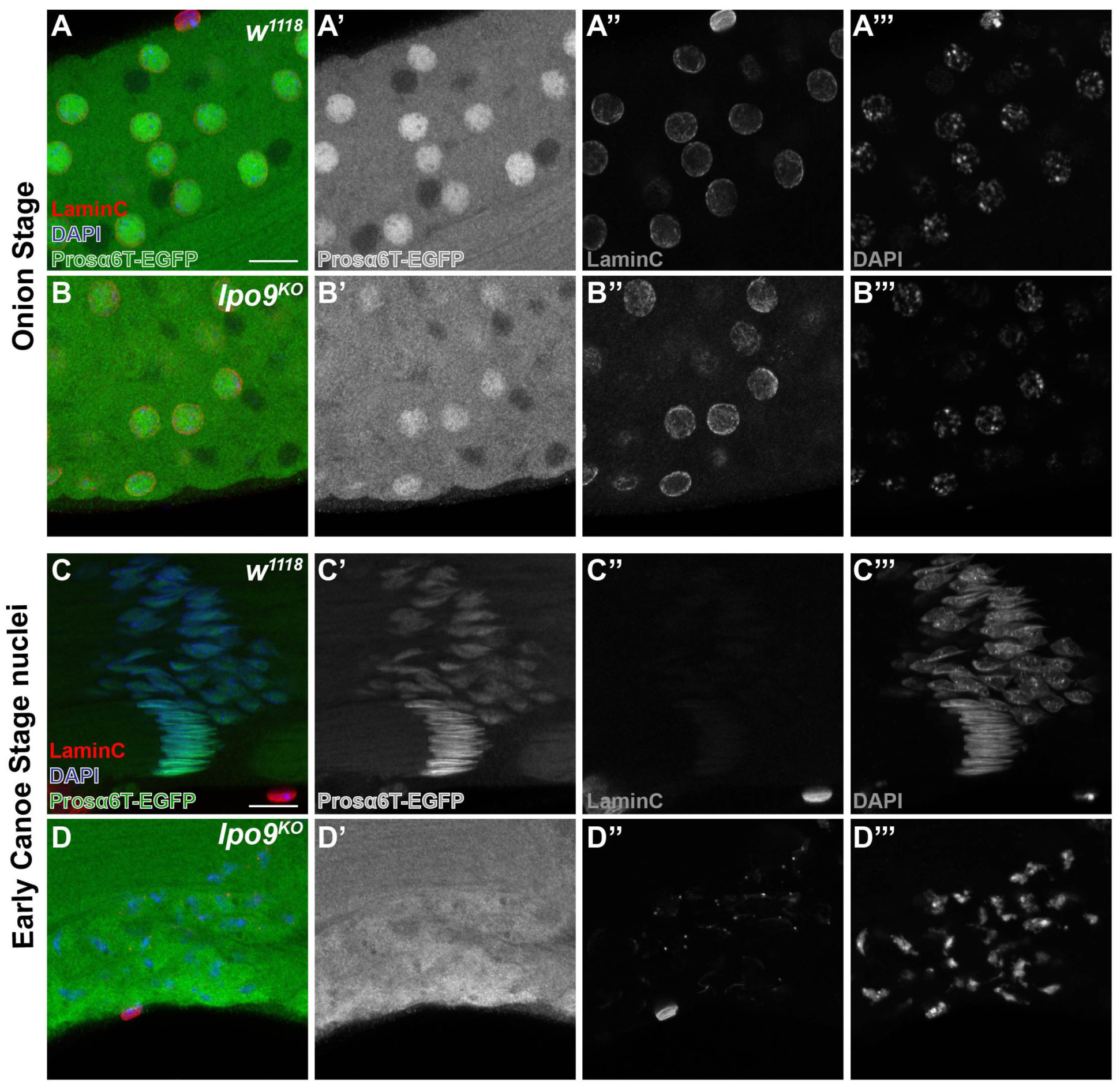
*Ipo9^KO^* spermatids show reduction of Prosα6T in the nucleus. (A-D) Spermatids at the onion stage and early canoe stage, stained for Prosα6T-EGFP (green), LaminC (red) and DNA (blue). (A & C) *w^1118^* spermatids and (B & D) *Ipo9^KO^* spermatids. Scale bars 10μm.

